# Measuring forces and stresses *in situ* in living tissues

**DOI:** 10.1101/016394

**Authors:** “Forces in tissues” workshop participants

## Abstract

Development, homeostasis and regeneration of tissues result from the interaction of genetics and mechanics. Kinematics and rheology are two main classes of measurements respectively providing deformations and mechanical properties of a material. They are now applied to living tissues and have contributed to the better understanding of their mechanics. Due to the complexity of living tissues, however, a third class of mechanical measurements, that of in situ forces and stresses, appears to be increasingly important to elaborate realistic models of tissue mechanics. We review here several emerging techniques of this class, their fields of applications, their advantages and limitations, and their validations. We argue that they will strongly impact on our understanding of developmental biology in the near future.

## I Introduction

During multicellular development, homeostasis and regeneration, tissues undergo extensive morphogenesis based on coordinated changes of cell size, shape and position over time. Such processes are under the regulation of genetics and mechanics. Genetic information is translated into biochemical signals, which regulate body axis specification, tissue patterning, sequence and timing of cell activities such as proliferation, migration, and differentiation. Mechanics provides another layer of regulations of development, which could be as evolutionally conserved as is genetic regulation (Savin et al., 2011). Mechanical forces and **stresses** (for terms in bold, see Glossary and Fig. 1) arise at the molecular level in part from the distributions and regulation of molecular motors (Lecuit et al., 2011). Cells integrate those forces to pull and push on their neighbors and ultimately deform tissues, depending on cell and tissue material properties (Heisenberg and Bellaïche, 2013; Sampathkumar et al., 2014). Forces and stiffnesses also control transcription programs of cell differentiation (Mammoto et al., 2012), especially in organs with a mechanical role (such as bones, heart, vessels or muscles). Forces determine cell changes during tissue repair and homeostasis (Guillot, 2013). Understanding force generation and sensation, feedback, and integration of genetics and mechanics in multicellular tissues is an active field of research which might lead to a comprehensive multiscale picture of tissue and organ development, homeostasis, regeneration, or disease.

**Figure 1.**
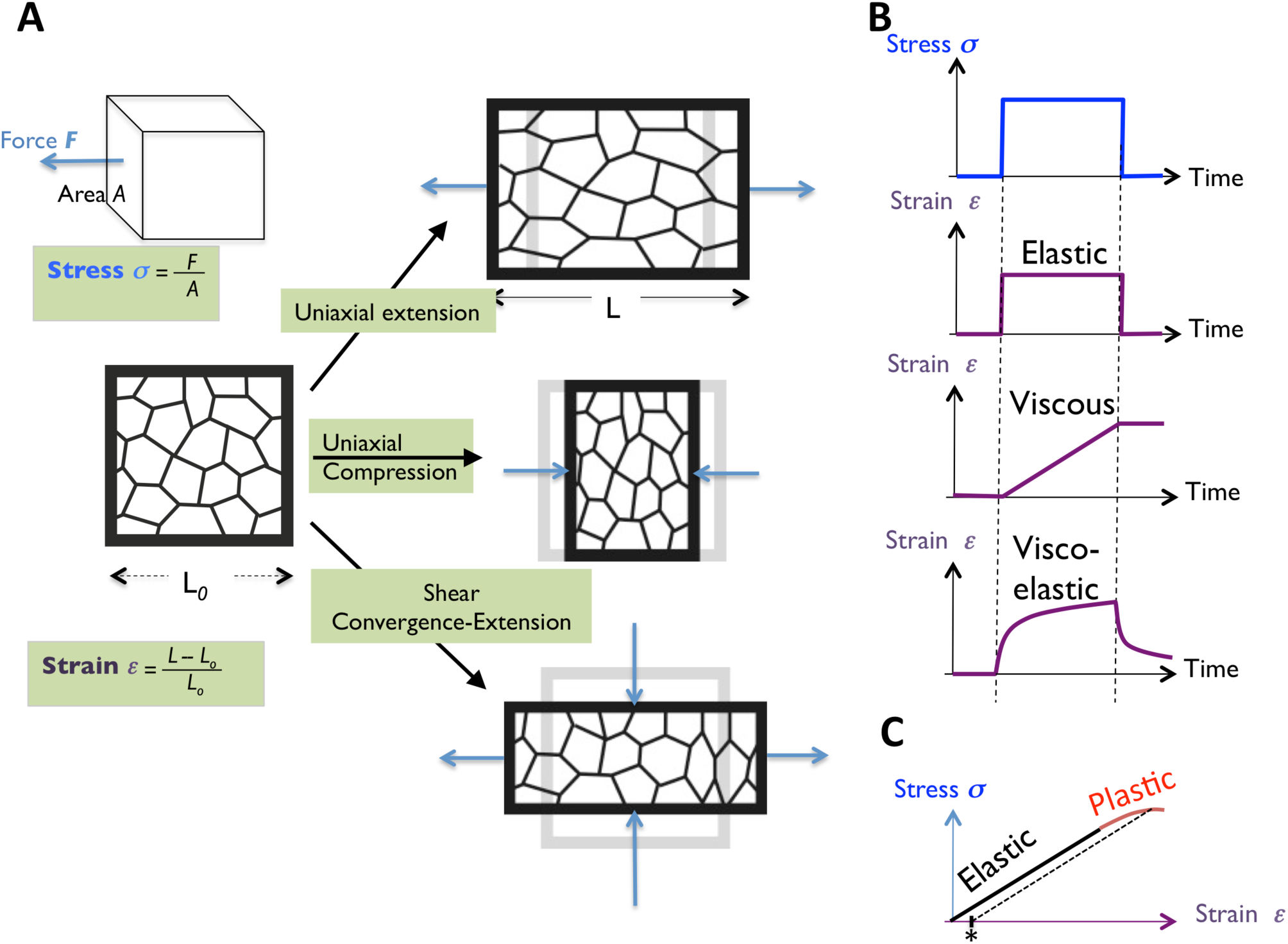
**Definition of mechanical terms.** (A) Schematics showing different types of deformation: uniaxial extension, uniaxial compression and pure shear (convergent-extension). When a material experiences a force perpendicular to one of its surfaces, stress is defined as the ratio of the force to the surface area *A*. Force has a magnitude and a direction; tension and compression correspond to forces pointing respectively outwards and inwards of the body. Strain is commonly defined as the fractional increment in length due to tensile or compressive stress. Shear (pure shear) corresponds to a constant volume strain, whereby the body is compressed in one direction and extended in the orthogonal direction. (B) Mechanical response (strain) of an elastic, viscous and visco-elastic material to the application of a constant stress and after the release thereof. (C) Stress-strain curve of an idealized material exhibiting elastic and plastic behaviors. The transition point is called the yield. In the plastic regime there is irreversible strain, which results in a residual strain. The residual strain (marked by an asterisk) induced by a given stress can be determined by a dashed line, drawn back to the strain axis with a slope equal to that of the initial elastic loading line.

In order to achieve this goal, quantitative characterisation of tissue mechanics is essential. Two classes of mechanical measurements have now been widely applied to cells and tissues: **kinematics** and **rheology** (Fung, 1993; Oates et al., 2009; Labouesse, 2011). Kinematics, usually provided by *in situ* live imaging of structures in cells and tissues, comprises the measure of structure (shapes, sizes and positions) and changes thereof in space and time, that is, **deformations** and **deformation rates**. Rheology comprises the measure of **viscosity**, **stiffness** and **yielding**, which are the respective mechanical properties that characterize a **liquid**, an **elastic solid** and a **plastic solid**. The third type of independent measurable variable regard **dynamics**, namely forces and stresses through which the tissue constituents interact mechanically.

In simple non-living materials, measuring mechanical properties and deformations allows to infer the involved forces and stresses, based on simple models and assumptions. While this approach may be applied to some extent to living materials, cells and tissues are typically complex systems with multiple mechanical structures intertwined over a wide range of scales that all generate forces, exhibit deformations and bear specific mechanical properties. As a consequence, development of more complex models, solving methods and assumptions are required to infer forces. In addition, deformations and mechanical properties may not be experimentally accessible at all scales. Hence, direct *in situ* measurements of forces and stresses in live cells and tissues appears as a valuable addition to establish sound, multiscale models of cell and tissue mechanics.

Measurements of force and stress have been up to now mainly performed in cells *in vitro* (Addae-Mensah, 2008). Yet, there is an emerging body of literature on methods (and applications thereof), either transferred from *in vitro* to *in vivo*, or recently developed for *in vivo.* We review here (Figs. 2-5) techniques which might become part of a versatile toolbox to study tissue dynamics *in vivo*, especially during development: first, contact methods; then, non-contact methods based on manipulation, sensors or imaging. For each method we explain which biological questions it can help to address, list which quantities it measures and over which range, discuss its advantages/disadvantages, and explicit the hypotheses it relies on. The cost of each technique is lower than that of microscopes: it is listed in Table 1, as well as scales, advantages and drawbacks. We end with some perspectives. We hope to help newcomers in the field to select techniques suitable to their questions, encourage the formation of a community with common cross-validation methods, and improve the integration of mechanics data into models to deepen our understanding of physical principles of development.

**Table 1.** Synthetic view of method comparisons.

|  | Measured quantities | Measurable range | Time scale* | Size scale ** | Advantages | Disadvantages | Cost |
| --- | --- | --- | --- | --- | --- | --- | --- |
| **Optical/magnetic tweezers** | Cell junction tension | pN-nN | ms-min | 0.1 – 10 $\mu\text{m}$ | Non-contact<br>Absolute measurements | Delicate calibration | €€€ |
| **FRET force probe** | Intramolecular tension | pN | >s | nm | Molecular measurements | Requires different control constructs; delicate calibration | € |
| **Birefringence** | Tissue-scale stress | >10kPa | >s | > $\mu\text{m}$ | Global | Required flat, transparent sample; delicate calibration | € |
| **Liquid drops** | Cell-scale stress | 4- 30 mN/m ;<br>0.3-60 kPa | 0.1 s-hours | > 5 $\mu\text{m}$ | Absolute measurements | Requires surface chemistry of droplets | €€ |
| **Indentation/ Microplates/ AFM** | Cell or aggregate surface tension | 0.1 Pa | s-hours | Few to 100 $\mu\text{m}$ | Absolute measurements | Contact method | €€-€€€€ |
| **Pipette aspiration** | Cell or aggregate surface tension | $\mu\text{N/m}$ -<br>mN/m | >10s | Few to 100 $\mu\text{m}$ | Absolute measurements | Contact method | €€ |
| **Subcellular laser ablation** | Cell junction tension to dissipation ratio | NA | s-min | 0.1 – 10 $\mu\text{m}$ | Non-contact | Possible collateral damage | €€-€€€ |
| **Tissue-scale laser ablation** | Tissue stress to viscosity ratio | NA | s-min | 10 $\mu\text{m}$ - 1 mm | Non-contact | Requires sample and laser alignment; one experiment per sample | €€-€€€ |
| **Force inference** | Relative cell junction tension, cell pressure | NA | video rate | > $\mu\text{m}$ | Image based, global | Requires image segmentation | € |

\* Time scale of the mechanical processes that can be probed
\*\* Size scale of the mechanical processes or mechanical elements that can be probed
Cost: € (< 10 k€); €€ (10-50k€); €€€ (50-100k€); €€€€(100k€-200 k€) excluding the microscopes
NA: not applicable since only relative measurements

**Figure 2.**
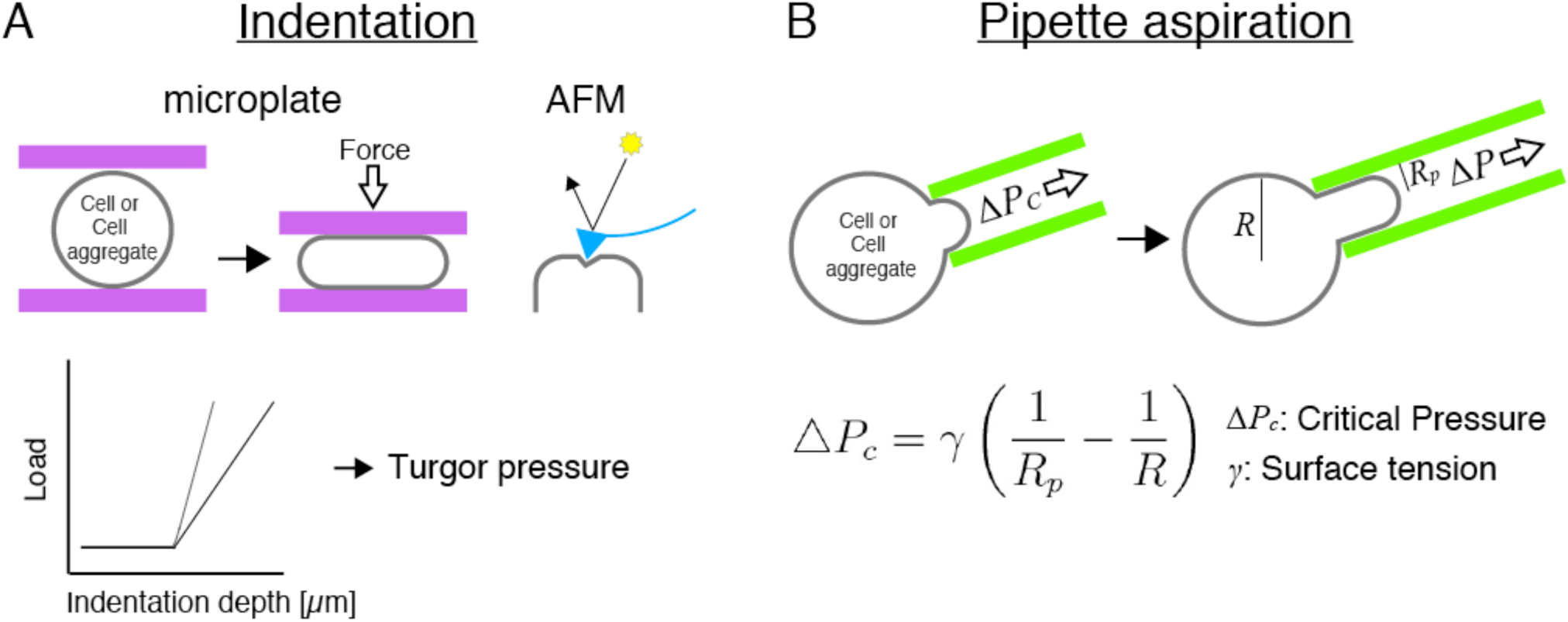
**Contact methods.** Micromanipulation by indentation (A) or aspiration (B). See text for details. Adapted with permission from: (B) Guevorkian et al., 2010.

## II Methods

### 1. Contact methods

We start with methods to infer forces/stresses (and sometimes also material mechanical properties), which involve a direct contact between the sample and the measurement apparatus: micromanipulation using microplates, atomic force microscopy, indenters and pipettes. The measurement apparatus either pushes (1a) (Fig. 2A) or pulls (1b) (Fig. 2B) samples.

#### 1a- Micro-manipulation: pushing

During morphogenesis, tissues change their shapes according to their mechanical properties and to internal and external forces. Micro-manipulation approaches aim at probing material properties and eventually internal forces by making mechanical contacts with the sample, by applying controlled external forces and by measuring deformation (Fig. 2A).

Micro-manipulation can be performed by tools such as microplates or indenters (Davidson and Keller, 2007), which provide information at different spatial scales. Microplates are parallel plates, one of which being a force transducer, which apply uniaxial compression on a large surface area of the sample causing the sample undergo strain, with for instance less than microNewton forces in the case of soft embryonic tissues (Foty et al., 1994; Fig 2A top left). Indenters push on the sample across a small contact area as compared to the whole tissue (Lomakina et al., 2004; Fig. 2A top right). There is a variety of indenters, ranging from atomic force microscopes (AFMs) capable of applying picoNewton to nanoNewton forces, to microscale indentors (often rod shaped), capable of applying microNewton loads: together, these tools cover six decades in forces.

A strong advantage of such approaches is that they give access to absolute values of surface **tension** (Table 1). Yet, they require direct contact to the sample; they apply to single cells or to tissues, which are directly accessible, without rigid external layer.

The main application of indenters, including AFM, is measurements of tissue explant stiffness (Mathur et al., 2001). AFM has been also used to infer contractility of the cortex (Krieg et al., 2008) from its effect on surface tension, which can be deduced from the cell's deformation in response to a controlled force. Differential tension probed by AFM functions to sort progenitor cells, leading to germ-layer organization of Zebrafish embryo (Krieg et al., 2008). Indenters have also measured turgor pressure in plants, whose contribution to the cell wall resistance can be explicitly separated from the cell wall visco-elastic properties (Forouzesh et al., 2013; Beauzamy et al., 2014). Developmental changes in turgor pressure proved to be finely tuned to support flower opening, anther dehiscence and lateral root emergence (Beauzamy et al., 2014).

#### 1b- Micro-manipulation: pulling

In micropipettes, an aspiration causes the deformation of the sample across the aperture of the pipette. The aspiration pressure is tuned at the critical value at which the cell deforms in the pipette to form a hemisphere with a radius equal to the radius of the pipette (Fig. 2B left). The equilibrium shape of the cell into a pipette results from the balance of its surface tension, to which contributes its contractile cortex (Evans and Yeung, 1989; Tinevez et al., 2009), and the pressure aspiration. The cell surface tension can then be simply determined from the pressure in the micropipette and the radii of the cell inside and outside the micropipette (Young-Laplace law): typical values range between tens of μN/m and tens of mN/m.

Enlarging the scale, pipette aspiration has been used to measure the surface tension of a group of sea urchin cells (Mitchison and Swann, 1954) and of mouse eggs (Larson et al., 2010). Such surface tension measurements provide absolute values with precision of a few percents: the biological variability dominates the experimental uncertainty. They require a few minutes and are suitable for morphogenetic events much slower than minutes. Surface tensions between tens of μN/m and tens of mN/m have been measured on cell aggregates (Norotte et al., 2008; Mgharbel et al., 2009; Guevorkian et al., 2010). With applied forces ranging between 0.1-10 μN, elastic moduli of few hundreds of Pa and viscosities of hundreds of kPa.s have been measured on cell aggregates.

Dual pipette aspiration has been recently implemented to quantitatively map all surface tensions of a mouse embryo in space and time (Maître et al, 2015). This study revealed that the first morphogenetic movement of the embryo, compaction, is mediated by increased acto-myosin contractility, rather than by increased cell-cell adhesion as commonly thought. This paves the way for quantitative mechanical studies of mouse embryogenesis.

### 2. Non-contact methods: manipulation

The use of contact methods is limited to applications in which direct access to tissues is possible. Yet, it is often required to implement non-contact methods to probe cell mechanics inside the organisms. In this section, we present optical or magnetic “tweezers”, which enable non-contact manipulation of objects such as microbeads inserted in the sample, to quantify forces (2a) (Fig. 3A, B). We then describe laser ablation, which also measures forces and stresses through non-contact manipulation of biological structures (2b) (Fig. 3C, D).

**Figure 3.**
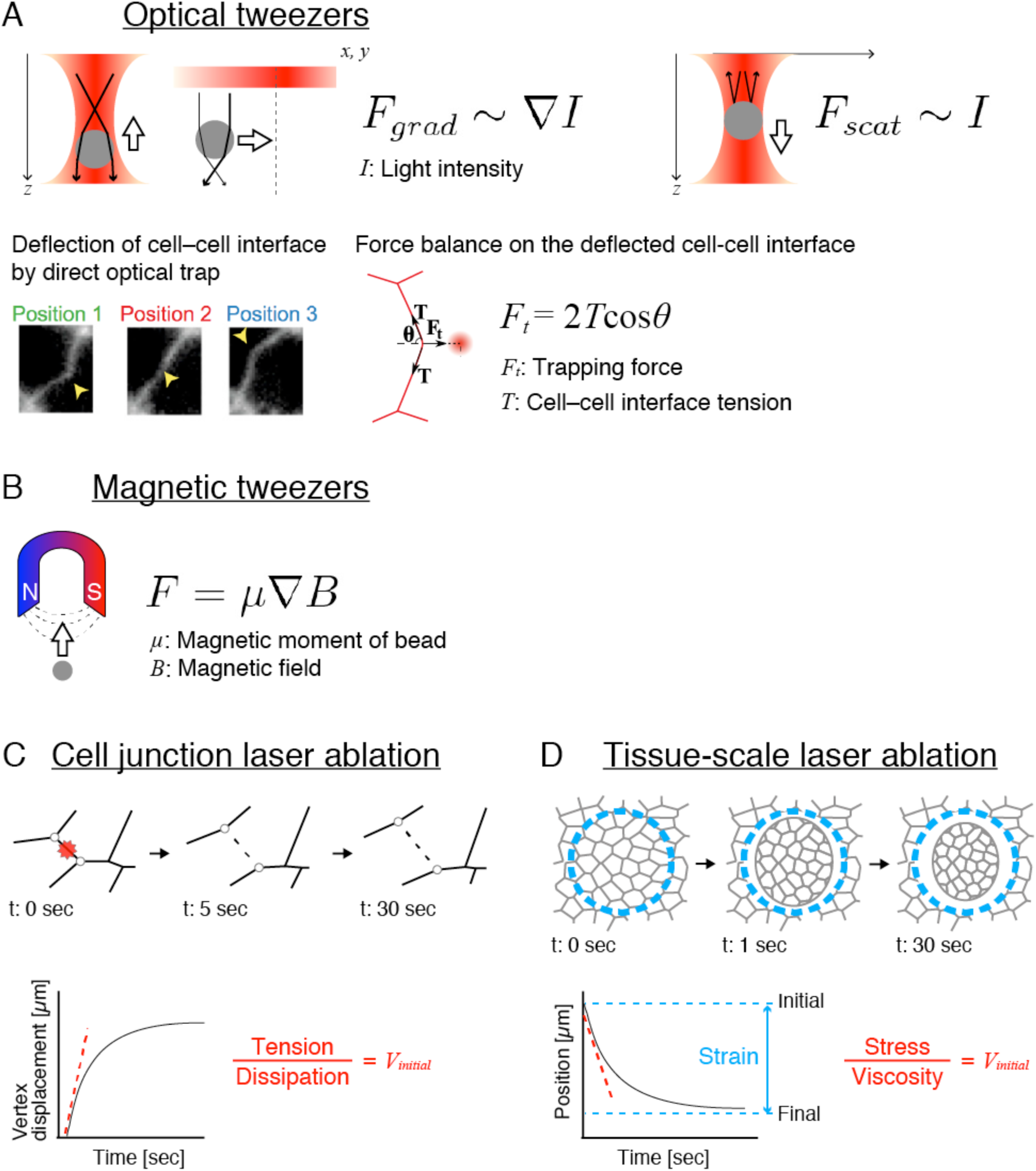
**Non-contact methods: manipulation** (A) Optical tweezers. (B) Magnetic tweezers. (C) Subcellular laser ablation. (D) Tissue scale laser ablation. See text for details. Adapted with permission from: (A) Bambardekar et al., 2015, (D) Bonnet et al., 2012.

#### 2a- Tweezers: optical, magnetic

Optical or magnetic tweezers refer to set-ups that apply forces to transparent or magnetic objects through gradients of light intensity or magnetic field, respectively (Fig. 3A, B). The applied forces are used to measure subcellular forces and cell mechanical material properties. The methods are generally semi-invasive, as they allow micromanipulation at a distance of a probe (transparent/magnetic bead or ferrofluid) in contact or within the biological sample.

The working principle of optical tweezers is that a transparent particle (showing a higher refraction index than the surrounding) in a focused beam is subject to a gradient force pulling it toward the high intensity region (focus) of the beam, while a scattering force pushes it away from the focus in the direction of incident light (Fig. 3A top) (Svoboda and Block, 1994). An effective optical trap occurs when the gradient forces dominate over the scattering forces. The optical force is a function of the geometry and refractive properties of the particle as well as of the intensity profile of the optical field. Optical tweezers can be combined to most microscope setups when the infrared laser used to produce the laser trap is focused by the same objective lens as the one used for image collection. As some organelles show refraction index mismatch with the surrounding, optical tweezers can trap them directly without the use of a particle, and the method becomes non-invasive (Bambardekar et al., 2015; Fig. 3A bottom).

The working principle of magnetic tweezers is that a magnetic particle in a magnetic gradient is subject to a force directed toward the source of the field (Fig. 3B). The force depends on the magnetic properties and geometry of the particle used as well as on the intensity profile of the external field. Magnetic fields are produced either by permanent magnets or electromagnets, which are mounted on a microscope stage and can come in different designs depending on the specific application pursued (Tanase et al., 2007).

Despite their potential advantages with regard to contact techniques, tweezers experiments have up to now mostly been used for cell rheology. In this case, the displacement of the tweezed object indicates the nature and the magnitude of the mechanical resistance, which opposes the experimentally applied force; the resistance can be elastic, viscous or a combination of both properties (viscoelastic material). Tweezers experiments can also yield access to quantitative force measurements provided *in situ* calibration, which might turn out to be delicate in living tissues. Measurable forces are typically on the order of pico- to nanoNewtons with magnetic tweezers and on the order of 1-100 picoNewtons with optical tweezers: together, these tools cover three decades in forces.

Recently optical tweezers have been applied in the early *Drosophila* embryo to measure forces at junctions; without even injecting beads, the same measurement has been performed by direct trapping of cell-cell interfaces (Bambardekar et al., 2015). This work revealed that tension at cell junctions is on the order of 100 pN and that tension equilibrates over a few seconds, a short timescale compared with the contractile events that drive morphogenetic movements, and delineated how local, subcellular deformations propagate in the plane of the tissue.

Magnetic tweezers have been used in *Drosophila* to produce tissue scale deformation thanks to the injection of large volumes of ferrofluid in the early embryo: magnetized cells were manipulated to impose compression, suggesting that germband extension upregulates Twist expression (Desprat et al., 2008).

#### 2b- Laser ablation: sub-cellular scale, tissue scale

Laser ablation experiments consist in severing biological structures taking part in the force transmission across the tissue (cytoskeletal filaments, cell-cell contacts). Assuming the tissue is close to mechanical equilibrium before ablation implies that forces conveyed by these structures almost perfectly balance each other. The severed force and the non-severed one are thus equal in amplitude and opposed in sign. Ablation provokes a sudden force imbalance (Fig. 3C, D): the dynamics of relaxation after ablation is governed by the non-severed force and the friction forces (proportional to the velocity which appears). The observed speed of relaxation of the structures just after ablation is thus a local measure of a ratio between the non-severed force and dissipation proportional to the material viscosity, hence indirectly measures the severed force to viscosity ratio.

Molecular bond rupture and laser-induced plasma formation (by generation of free electrons) are the main processes underlying laser ablation. Their production depends on different parameters such as laser wavelength, intensity, focal volume, pulse duration and pulse repetition rate (Vogel and Venugopalan, 2003). In practice two types of laser are used in living tissues. Ultra-violet nanosecond pulsed laser is cheaper, and it is easier to set a suitable light intensity. Near-infrared femtosecond pulsed laser (NIR-fs) offers the advantage to produce less collateral damages and has been shown to preserve membrane integrity, while allowing disruption of cortical cytoskeleton (Rauzi et al., 2008).

In subcellular ablation, a tightly focused laser is targeted at cell-cell junctions to disrupt the biological structures which support tensile forces (such as the actomyosin network). This results in a force imbalance, which drives changes in cellular geometries (cell vertex displacement for instance), usually visualized with fluorescent reporters of the structures of interest. It measures the ratio of junction tension to dissipation. It can be used as a proxy for relative value measurements of junction tensions (Fig. 3C) when one can assume that the dissipation changes much less than the tension between compared conditions.

The advantages of subcellular laser ablation are that it is a non-contact method, and it is easily combined with different types of microscopes using simple optical elements. These advantages allow its application to a large variety of tissues in organisms such as *Drosophila* (Farhadifar et al., 2007; Kiehart et al., 2000; Rauzi et al., 2008) or C. elegans (Mayer et al., 2010). Relaxation measurements after ablation require a few minutes and can be repeated few times in the same sample (at different positions in the embryo). In these different systems, laser ablation was key to reveal how cell contractility drives morphogenetic movements. In *Drosophila*, laser ablation revealed that anisotropy of tension at cell junctions drives cell intercalation and tissue elongation (Rauzi et al., 2008). In the C. elegans single cell embryo, anisotropy of cortical tension measured by laser ablation were shown to produce cortical flows, which establish antero-posterior polarity (Mayer et al., 2010).

Enlarging the scale to directly probe the stress at the tissue level, a laser can simultaneously ablate several cells (Hutson et al., 2003; Behrndt et al., 2012; Campinho et al., 2013). The initial velocity after ablation measures the stress-to-viscosity ratio within the tissue, in the direction of the relaxation. Alternatively, severing a large circle (Bonnet et al., 2012) reveals the main directions of stress and its anisotropy (Fig. 3D). In addition, having isolated a piece of tissue reveals its final relaxed stated, yielding the strain along different orientations with percent accuracy. With additional hypotheses and a model, information can be obtained about the ratio of external friction to internal tissue viscosity.

In *Drosophila*, large scale laser ablation revealed that dorsal closure is mechanically driven by the contractility of a supracellular actomyosin string and amnioserosa cells (Hutson et al., 2003). It was also implemented to understand how planar cell polarity can control mechanics (Bonnet et al., 2012). The ablation time and the time between successive images should be less than a second to remain much shorter than the relaxation time. The maximal ablation scale is determined by the tissue curvature, since all cells to be ablated should be simultaneously in focus. Proper axial and lateral alignment is required for efficient and well-controlled ablation experiments, and becomes crucial for the large scale circular ablation.

At even larger scale (millimeters), the same principle applies when cutting with a scalpel a tissue under tension (Beloussov et al., 1975). In flat tissues such as a floating biofilm, absolute values in the milliNewtons are measured up to percent precision in an inexpensive set-up using a standard force sensor (Trejo et al., 2013).

### 3. Non-contact methods: visual, with sensors

We next present visual methods which estimate forces or stresses by measuring passive changes of probes at molecular scale (FRET tension sensors, 3a) (Fig. 4A) or cellular scale (liquid droplets, 3b) (Fig. 4B).

**Figure 4.**
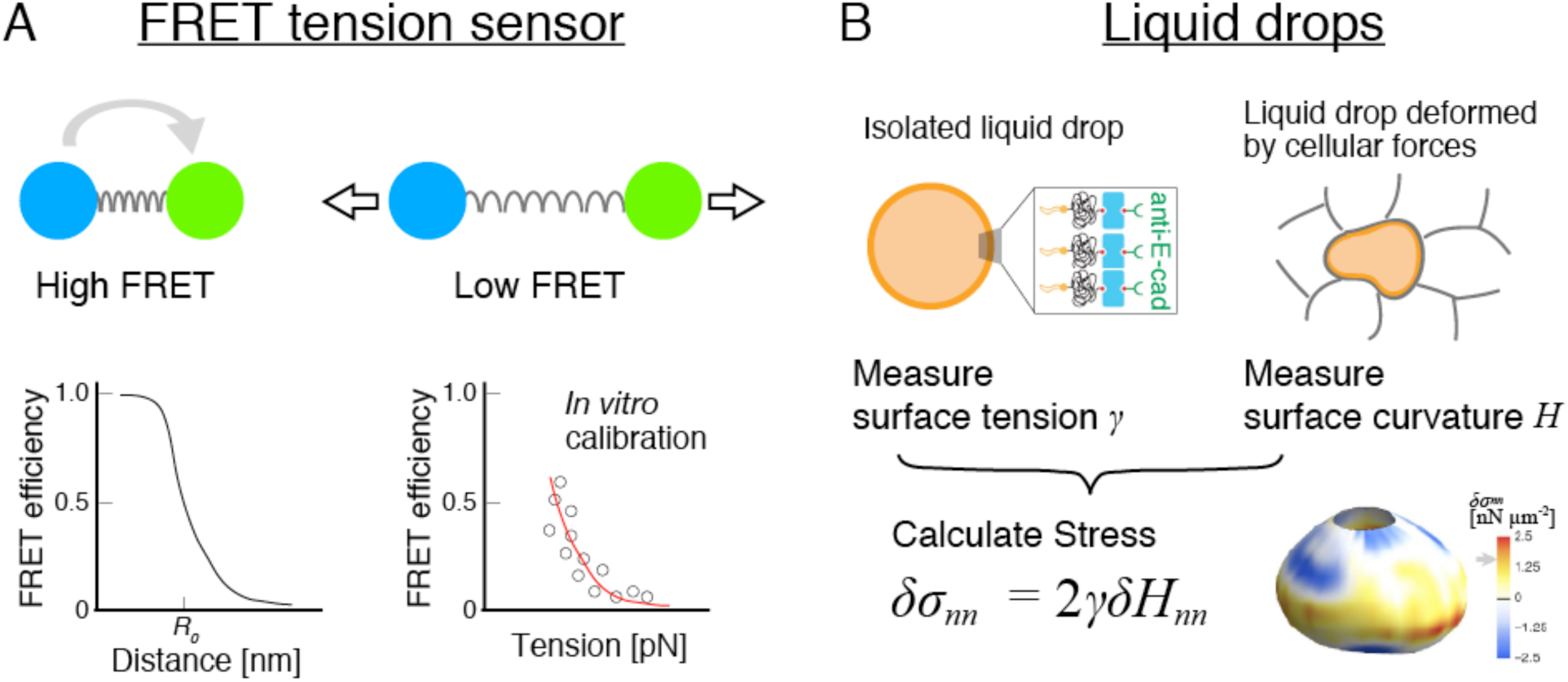
**Non-contact methods: visual, with sensors.** (A) FRET tension sensor. (B) Liquid drop stress sensor. See text for details. Adaptated with permission from: (A) Grashoff et al., 2011 and (B) Campàs et al., 2014.

#### 3a- FRET-based molecular tension biosensors

Force is supported by molecular elements such as cytoskeletal and adhesion components, which are capable of producing and/or transmitting forces. Förster resonance energy transfer (FRET) can be used to measure the tension amplitude (but not direction) that specifically these molecules experience in tissues.

FRET is an energy transfer between transition dipoles of two fluorophores (Fig. 4A), namely donor and acceptor (Miyawaki, 2011). Its efficiency *E* sharply depends on the distance *R* between fluorophores, as a power 6: *E = R_0_^6^/R^6^ + R_0_^6^*, where *R*_0_ is the distance where FRET efficiency is 50 % (Fig. 4A bottom left). The measured value of its efficiency is highly sensitive to variations in *R*, hence its nickname “spectroscopic ruler”.

Now, if a spring of known stiffness is inserted between donor and acceptor fluorophores, the change in spring length measured by FRET enables to infer the tension in the spring. A tension sensor such as a polypeptidic elastic spring flanked by two fluorescent proteins suitable for FRET is genetically encoded and inserted into a protein of interest by using standard molecular biology techniques to report the tension in that protein (Meng et al., 2008; Grashoff et al., 2010; Fig. 4A top). Truncating or mutating binding sites within the protein of interest provides variants that may be used as tensionless controls. FRET changes can be measured in fluorescence microscopy from changes in donor fluorescence lifetime, or relative changes in donor and acceptor intensities. Whether fluorescence lifetimes or intensities are measured, calibration measurements usually using FRET control constructs are required to compute changes in FRET efficiency. Then, if using a polypeptidic spring which FRET efficiency-force relationship was previously calibrated by single-molecule tweezing *in vitro* (Grashoff et al., 2010), molecular tension can be quantitatively obtained. Such a spring was designed to probe the 1 to several pN force range (Fig. 4A bottom right). The measured tension is a scalar, and an ensemble measurement on all the proteins comprised in the optical volume probed (which size is at least that of the focal volume), within the acquisition time, typically 100 ms.

A major interest of the method is that it could be applicable to any protein of interest as long as the tagged protein retains the functions of the native protein. This can be tested at the molecular, cellular, and organism level using various assays, to the extent of the prior knowledge of the protein functions. In addition, the method is in principle compatible with imaging of other proteins and/or structures, other force measurements above and below it, or any mechanical manipulation performed on a microscope stage. The method relies on molecular tension being the only cause of FRET change, which can be tested using tensionless control constructs under the same perturbations as their tension-sensing counterpart. In principle, non-tension contributions to FRET changes, regardless of their origin, may, this way, be corrected for. The method assumes that inserting a tension sensor module into a host protein does not change the efficiency-force relation, and that the calibration *in vitro* remains valid *in vivo*. Finally, the method does not discriminate molecular tensions at and above its detection range upper limit, thus possibly underestimates molecular tensions.

FRET tension sensors have been successfully applied to adhesion and cytoskeleton-related proteins in cell culture and *in vivo* model systems (Grashoff et al., 2010; Borghi et al., 2012; Krieg et al., 2014; Cai et al., 2014). In *C. elegans*, the method revealed that tension in β-spectrin ensures the structural integrity of touch receptor neurons and supports efficient sensation of touch (Krieg et al., 2014). The tension sensor has also been recently shown to behave as a uniaxial compression sensor in cell culture (Paszek et al., 2014).

More generally, fluorescent materials exhibit a variety of properties that could be dependent on mechanics (Gomez-Martinez et al., 2013; Watanabe et al., 2013). This opens up an opportunity to design and develop new force probes in the near future.

#### 3b- Liquid drops

Liquid drops can be used as versatile stress sensors both *in vivo* and *in vitro* (Campàs et al., 2014) (Fig. 4B). In the absence of externally applied stresses, the drop is spherical because of its surface tension. When a drop is introduced between cells in a tissue, it is deformed by local stresses. Confocal microscopy is used to determine the drop shape in 3D. Provided that the drop interfacial tension is separately measured, analysis of the droplet shape directly provides a measure of the anisotropic stresses locally exerted on the drop by the neighboring cells or the extracellular matrix (equation in Fig. 4B).

Producing the drops requires basic chemistry, which costs more in term of work than of reagents (typically hundreds of euros). Drops used in stress measurements should be made of bio-compatible oil (e.g., fluorocarbon oil), fluorescently labelled, cell-sized and coated with ligands for cell adhesion receptors, so that both tensile and compressive stresses can be measured. The equipment includes a fluorescent microscope, preferably with the ability to perform 3D imaging (such as confocal or two-photon microscopes); any apparatus classically used to measure surface tensions of liquids; and a micro-injector. A detailed description of how to make and use liquid drops as stress sensors can be found in (Paluch, 2015; chapter 20).

The experimentally obtained drop shape is analysed with nearly pixel resolution with a homemade software to obtain the local curvatures and, thereby, the cellular stresses on the droplet surface. The experiment can last for days without affecting normal embryonic development, provided the drops are small enough compared to tissue size. The drop size cannot exceed the capillary length (600 micrometers for fluorocarbon oil), as the drop shape would otherwise be affected by gravity. Droplet sizes larger than cell size (typically about 10 micrometers in diameter) are required to avoid being internalized by cells.

Drop contour curvatures can be measured typically with 5% accuracy (this value depends on image resolution and the value of the curvature itself), so that the limiting factor is the precision of interfacial tension measurement, around 20% for the minimal interfacial tensions used (approx. 3 mN/m). This relies on the hypothesis that the interfacial tension remains constant and uniform when the drop is introduced between cells. Such hypothesis is reasonable if the drop surface has been beforehand saturated with surfactants, but should be validated independently. The value of the interfacial tension used can be tuned to enlarge the range of measurable stresses, from approximately 0.1 to 60 kPa.

This method has already been used to prove that anisotropic stresses generated by mammary epithelial cells are dependent on myosin II activity (Campàs et al., 2014). In situations where the drop is only partially embedded in the tissue and displays a free surface, it is possible to measure the absolute value of the stresses and not only the anisotropic stress.

Although cells may react on their neighbours' stiffness, experimental data indicates that mesenchymal cells of mouse embryonic tooth can generate the same stresses in aggregates and in living tissue, for both drops with small and large interfacial tension, lending confidence to the method's validity and reproducibility. Additionally, cell disruption using detergents (SDS), induces the immediate rounding of the drops towards a spherical shape.

A possible variant would consist in replacing the liquid drop with a soft spherical elastic bead. If deformations remain small, fitting an ellipsoid to the bead contour would yield the main directions of stress applied by cells on the bead. The change in bead volume would measure the pressure or traction in the surrounding tissue. The beads could be made of polymers in a microfluidic setup; their stiffness should be adjusted, typically around 1 kPa, and carefully calibrated. Alternatively, gel sensors can be external instead of internal: embedding a Xenopus embryo in a soft agarose gel has already enabled to estimate the forces it develops during convergent extension (Zhou et al., 2015).

### 4. Non-contact methods: purely visual

We finally present purely visual methods, which use either intrinsic material properties of the tissue (birefringence, 4a) (Fig. 5A, B) or cell shapes (force inference, 4b) (Fig. 5C).

**Figure 5.**
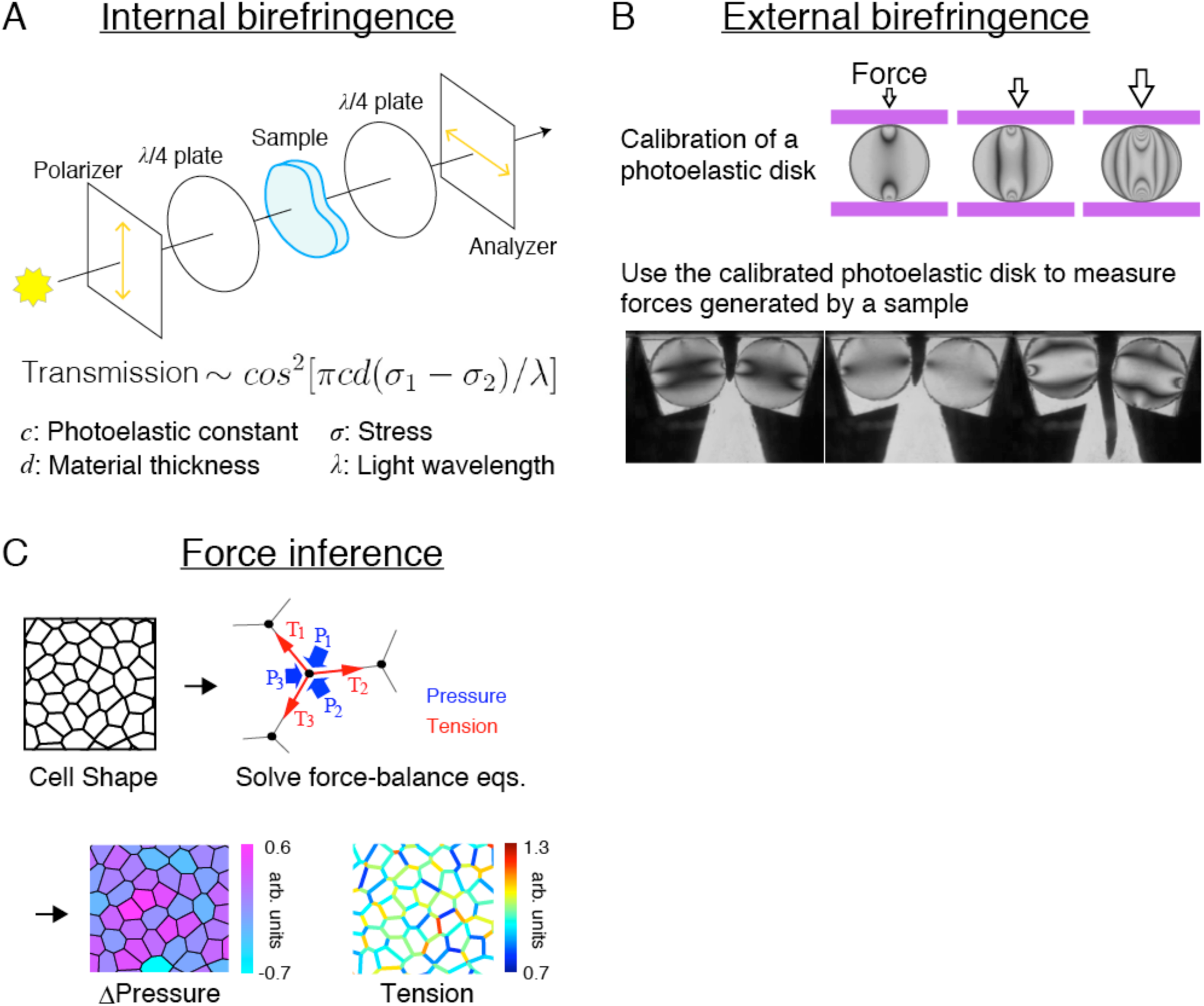
Non-contact methods: purely visual. (A) Internal birefringence. (B) External birefringence. (C) Force inference. See text for details. Adapted with permission from: (B) Kolb et al., 2012, (C) Ishihara and Sugimura, 2012.

#### 4a- Birefringence: internal, external

Tissue stress anisotropy, which results from the anisotropic spatial distribution of molecular and cellular forces, can be probed by tissue optical properties. The intensity of light transmitted by a flat object can be affected by its in-plane stress anisotropy. An image of a tissue obtained with a camera (with or without microscope, depending on scale, e.g. from micro- to millimeter) can hence be turned into a map of stress anisotropy within the tissue. The principle uses polarized light, as follows.

When an otherwise isotropic material is subject to a stress, its electronic structure differs in the directions of larger and smaller stresses. Such material, said to be “birefringent” (Fig. 5A), becomes optically anisotropic: its refractive indices in both directions *n*_1_, *n*_2_ differ by a term proportional (up to the so-called “photo-elastic constant” *c* which characterizes the material) to the stress differences along both axes. By placing it between two sets of combined linear and quarter-wavelength polarizers (Fig. 5A, top), the transmitted light intensity varies as a squared cosine of the index difference *n*_1_-*n*_2_ times the material thickness *d*.

The material thus acts as an indirect stress probe (Fig. 5A, bottom), which sensitivity is determined by both *c* and *d*. The object studied should be flat (or the profile of *d* is known), transparent and homogeneous. Measurements are non-destructive, and simple if stress is the only cause of anisotropy. The signal is integrated along the *z*-axis, so the information is mapped in the *xy* plane, and its space-time resolution is that of the camera. A crude version can be implemented on any microscope or with any camera for a few euros. The resolution can be improved by: using a liquid crystal retarder (for a few kEuro), optimizing image processing (Shribak and Oldenbourg, 2003), or using several polarizer and analyzer orientations which in addition determines the stress directions.

The relative precision can reach 10% while absolute calibrations require the determination of *c*, which is delicate *in vivo*. Within a tissue, *c* can be measured in an explant if birefringence measurements are combined with a micro-manipulation setup to apply a stress. In *Drosophila* wing discs (Nienhaus et al., 2009), but also in other tissues (plants, trachea) the photoelastic constants are of the order of 10^-10^ Pa^-1^ and stresses of 10 kPa are detectable (Schluck and Aegerter, 2010). Forces exerted by cells on a solid substrate down to 1 pN per square microns have been measured (Curtis et al., 2007).

The method has probed developmental changes in the distribution of stress in the *Drosophila* wing disc (Nienhaus et al., 2009). The stress distribution pattern supports theoretical models of wing disc growth (Hufnagel et al., 2007; Aegerter-Wilmsen et al., 2007). Future studies will combine the method with genetical and mechanical manipulations of a tissue and seek to clarify the mechanism, by which the reciprocal interaction between cell proliferation and tissue stress deforms a tissue into its final shape and size.

Alternatively, the birefringence of an external sensor placed in contact with the tissue yields a pattern of optical fringes related to the stress field in the sensor (Fig. 5B, top). This yields semi-quantitatively the force exerted by the tissue on the sensor (to make it quantitative requires to assume that the photoelastic material is truly linear, and to solve inverse problems which are sometimes difficult). It is biocompatible, non-destructive and suitable for living systems above a millimeter that move and/or grow. Validation is direct if the photoelastic material is well calibrated: either commercial (a few hundred euros for a sheet in which several sensors can be cut), or preliminary calibrated plexiglass or agar gels. This method was used to quantify radial force down to one Newton during root growth in response to constraints (Fig. 5B, bottom), which provided insights on mechanical interaction between the radial growth of roots and the porous or crack networks of soils and substrates (Kolb et al., 2012).

**4b- Force inference: static, dynamic**

In epithelial tissues where cell shapes are determined by cell junction tensions and cell pressures, i.e. assuming mechanical equilibrium, information on force balance can be inferred from image observation (Fig. 5C). For instance, if three cell junctions which have the same tension end at a common meeting point, their respective angles should be equal by symmetry, and thus be 120° each. Reciprocally, any observed deviation from 120° yields determinations of their ratios. The connectivity of the junction network adds redundancy in the system of equations, since the same cell junction tension plays a role at both ends of the junction.

Mathematically speaking, there is a set of linear equations, which involve all cell junction tensions, and cell pressure differences across junctions. The coefficients of these equations are determined from observation of vertex positions. The number of unknown can be larger or smaller than the number of equations, resulting in under- or over-determination, respectively. According to the mechanical and morphological properties of the system of interest, one of the following variants could be chosen.

Treating cell junctions as straight and neglecting pressure differences results in overdetermination (Chiou et al., 2012). Treating more general cases which are underdetermined, for instance when taking into account the pressure differences (assuming small cell curvatures), is possible using Bayesian force inference: it treats the ill-conditioned problem and simultaneously estimates tensions and pressures by using Bayesian statistics (Ishihara and Sugimura, 2012; Ishihara et al., 2013). Adding visual observation of cell junction curvature brings back to overdetermination, by including the Young-Laplace law that relates the cell junction curvature to tension and pressure (Brodland et al., 2014; Paluch, 2015, chapter 18). For more quickly evolving tissues, a dynamic version of force-inference called “video force microscopy” assumes the type of dissipation and uses temporal cell shape changes to infer forces. The original version solved an over-determined problem with a special three-dimensional geometry of cells (Brodland et al., 2010). A revised version is applicable to a general epithelial geometry (Brodland et al., 2014).

These variants of the inverse problem infer tension to only one unknown constant, the tension scale factor, whether for a few cells or for a few thousands. In the latter case, detection of contours of thousands of cells and precise measurement of the cell contact surface curvature requires a sophisticated image processing algorithm. When pressures are determined, there is an additional unknown constant, the average cell pressure. By integrating tensions and pressures, tissue stress can be calculated at all places, again up to these two constants. These constants are not visible on the image: if they are necessary, they have to be determined by another method.

The advantages of force inference are as follows. First, it offers both single cell resolution and a global stress map, which allows to compare the inferred forces directly with the activity of the molecules and with the cell-level dynamics, as well as with their effect on tissue scale. Second, it is non-destructive, so that the dynamics of forces or stresses can be analysed along a live imaging movie in two or three dimensions. Third, the method is free from any assumption regarding the biophysical origin of tension and pressures. Fourth, force inference lends it well to various validations, for instance against myosin distribution and cell junction ablation; tests on a controlled pattern prepared using numerical simulations have shown its accuracy and robustness to added noise (Ishihara and Sugimura, 2012). It is also suitable to perform cross-validations against other measurement methods.

Cross-validation has already begun in the case of Bayesian force inference. In *Drosophila* epithelial tissues, Bayesian force inference has been successfully compared to large-scale ablation anisotropy measurements (Ishihara et al., 2013) and photo-elasticity patterns (Ishihara and Sugimura, 2012). Force inference could itself become an absolute method if its unknown constants are determined independently, e.g. by optical tweezers, and subsequently serve as a common benchmark to validate other relative measurement methods.

Bayesian force inference reported the distributions of tension, pressure, and stress in the *Drosophila* pupal wing and showed that the extrinsic stretching force provides directional information for assignment of the orientation of cell rearrangement, leading to tissue elongation and hexagonal cell packing (Sugimura and Ishihara, 2013).

## III Concluding remarks

We have presented techniques at various scales, which provide the magnitude and distribution of forces and stresses shaping tissues during development. We now present perspectives: comparison and combinations of methods, and links between these experiments and models.

### - Combinations and comparisons

These techniques are diverse enough that they are complementary: for each system and biological question, one technique or the other should be more suitable. Altogether, they probe forces over a wide range of length and time scales. Importantly, most mechanical measurements are fast compared to the timescales of morphogenetic events from few tens to thousands of seconds (see Table 1) and are thus capable to follow the temporal changes of forces during development.

In practice, their fields of application are diverse too. Internal birefringence, optical tweezers and laser ablation require tissue transparency and mainly apply to thin or external tissues. Pipette aspiration and indentation act on cells accessible to direct mechanical contact. Drops and beads require injection. FRET, force inference and drops rely on fluorescence imaging, and apply to both two and three dimensions as deeply as the signal of the probe is detectable. Most methods rely on different assumptions.

It will be important to combine and apply these methods in a more integrated way. Some systems allow the simultaneous use of methods that measure different quantities, or at different scales or in different places. For instance, the combination of molecular-scale tension measurements (FRET sensors) with cell- and tissue-scale tension measurements or manipulations (e.g. with optical tweezers) may enable to propose models of force integration from the molecule to the cell, and thereby provide hints of the cellular mechanical architecture. Measuring forces is one contribution to our understanding of the link between the scales of intracellular components, cell, group of cells, and tissue-scale description in terms of stresses. Measurements obtained using a small scale probe can be gathered and presented over a much larger range. For instance, FRET-based molecular tension microscopy, force inference and birefringence are measured at the scale of molecules in a focal volume, cell junction and group of cells, respectively, but each can be suitably averaged over space to yield a tissue-scale map of some stress components. We also expect that combination of mechanical measurements with the fast-growing optogenetic tools will unveil important connections between mechanics and biochemical signalling.

Pairwise comparisons of two methods that measure the same quantities at the same scale in the same place allow cross-validation. Cross-validated results are less dependent on assumptions, and help a consistent picture emerge. Pairwise comparisons of some of these techniques are possible, and adequate systems can be designed to enable scale and penetration overlap, and calibration. Such systems include cultured cell monolayers, reconstituted cell aggregates, or even foams. In the future, establishing more *in vivo* systems enabling for cross-validations will speed up the development of *in vivo* force or stress measurement methods.

### - Links with modeling

Models based on mechanics are increasingly used in developmental biology (Keller, 2002). Whether using analytical equations or numerical simulations, they assist force and stress measurements in several respects (see Brodland, 2004; Anderson et al., 2007; Oates et al., 2009 for details of models used in tissue mechanics). For instance, theoretical reasoning helped to propose and design new force measurement techniques and experiments such as tissue-scale laser ablation, drops, or optical tweezers without beads (Bonnet et al., 2012; Campàs et al. 2014; Bambardekar et al., 2015). Models and assumptions are sometimes required to extract measurements of relevant parameters, if necessary through fits of models to data, notably in laser ablation and force inference (Hutson et al., 2003; Farhadifar et al., 2007; Ishihara and Sugimura, 2012; Brodland et al., 2010, 2014). Numerical simulations provide benchmark data in controlled conditions, to validate an experimental measurement, and test its sensitivity to a parameter or to errors (Landsberg et al., 2009; Ishihara et al., 2013; Brodland et al., 2014; Bambardekar et al., 2015). Finally, comparing experiments with analytical models or numerical simulations enable to refine the interpretation of experimental results and extract from them more information, such as material properties, or quantities which are not directly accessible to experiments (Farhadifar et al., 2007; Krieg et al., 2008; Rauzi et al., 2008; Mayer et al., 2010; Bonnet et al., 2012; Maître et al., 2012; Sugimura and Ishihara, 2013; Forouzesh et al., 2013).

While models help force and stress measurements, the opposite is true as well. Most models of tissue mechanics hypothesize material properties and distribution of forces, which are often not measured. Not only quantitative measurements will allow us restrict the number of unknown parameters but also help us making the right hypotheses, by testing falsifiable predictions of different models. Models could then confront images (or movies) and force measurements to determine the tissue material properties. This step is essential to develop predictive models of tissue mechanics, in an attempt to explain how cell-level changes (deformation, growth, displacement, neighbour swapping, divisions and apoptoses) do determine, though mechanical interactions, the final shape and size of a tissue (Tlili et al., 2015).

### - Future directions

Historically, due to practical and conceptual reasons, morphogenetic movements taking place at the same time in an organism have been studied separately one from another. Yet, with force and stresses measurements along with new imaging approaches, we expect a more integrated understanding of tissue interactions, tissue shape and movements. Force and stress measurements will be crucial to understand how the tissue mechanical properties are related one to another, from the molecule scale to the scale of the cell, tissue and whole organisms and from millisecond to hour timescales. It is likely that they will help us understand emergent properties of tissues, such as the simultaneously solid-like and fluid-like mechanical behaviours of tissues at the developmental timescales (hours). This dual mechanical behaviour arises from the relatively fast dynamics of cytoskeletal and adhesive structures, and cell mechanical activity due to molecular motors, which generate and transmit forces. Understanding this picture should illuminate the coupling between biochemical and mechanical regulations in tissues.

## Acknowledgments

The content of this article is the work of the participants to the informal and intense “Forces in tissues” workshop held in Paris 7 University, May 2014. The text has been written by Kaoru Sugimura, François Graner, Pierre-François Lenne, organisers of the workshop. For details and participants list, see http://www.msc.univ-paris-diderot.fr/tissue-stress.

## Glossary

Deformation: relative change in size of an object. In one dimension, it is a dimensionless number: the fraction of change in the object length, whether positive (elongation) or negative (contraction). In two or three dimensions, elongation in one direction can coexist with contraction in another direction, so that the deformation is described by amplitudes in different directions (see “dilation” and “shear”). Also called “strain”.
Deformation rate: deformation per unit time, expressed as the inverse of a time. In one dimension, it can be for instance a change of a few percent per hour. In two or three dimensions, it is more complex (see “deformation”), and is equal to the symmetrical part of the velocity gradient.
Dilation: component of the deformation, which is isotropic, corresponding to a change of size.
Dynamics: description of a material evolution in reference to the forces which cause it.
Elastic deformation: reversible part of the deformation.
Elastic modulus: see “stiffness”.
Elasticity: capacity of a material to deform and store energy in response to an applied stress, and reversibly return to its initial rest shape after stress is removed.
Kinematics: visual description of motion, without reference to force
Liquid-like behaviour: behaviour of a material with a non-defined rest state, which can be continuously deformed by a very small shear stress.
Plastic deformation: irreversible part of the deformation.
Plastic material: material that undergoes irreversible deformation in response to an applied stress.
Plasticity: capacity of a material to be sculpted, i.e. acquire a new rest shape under an applied stress, and keep it when the stress is removed.
Purely elastic material: Material that responses immediately and elastically to applied stress.
Purely viscous material: material that flows in response to a shear stress and does not recover its initial shape after stress is removed.
Rheology: science of flow and deformation of materials.
Shear: component of the deformation that is purely anisotropic, that corresponds to a change of shape.
Solid-like behaviour: behaviour of a material with a defined rest state.
Stiffness: quantity which measures the resistance of an elastic solid in response to an applied stress. Also called “elastic modulus” or “rigidity”.
Strain: see “deformation”.
Stress: here mechanical stress, i.e. forces internal to the tissue, exerted by a tissue region onto a neighbour tissue region. It is a more coarse-grained notion than force, and is expressed as a force per unit size of tissue boundary.
Tension: force internal to an object, tending to decrease its size. The “surface tension” of an aggregate or a piece of 3D tissue tends to decrease its surface area, and is expressed in Newtons per meter. A one-dimensional object like a cell-cell adherens junction has a “line tension”, tending to decrease its length, and expressed in Newtons.
Viscosity: (1) tendency of a material to resist gradual deformation and flow in response to an applied stress; (2) quantity which measures this property (see “viscous modulus”).
Viscous modulus: quantity which measures the resistance of a viscous liquid in response to an applied stress. Usually called “viscosity” for short.
Yield: here, defines the transition between the elastic behaviour and the plastic behaviour.

